# DeepCCI: a deep learning framework for identifying cell-cell interactions from single-cell RNA sequencing data

**DOI:** 10.1101/2022.11.11.516061

**Authors:** Wenyi Yang, Zhaochun Xu, Meng Luo, Yideng Cai, Chang Xu, Pingping Wang, Songren Wei, Guangfu Xue, Xiyun Jin, Rui Cheng, Jinhao Que, Wenyang Zhou, Fenglan Pang, Huan Nie, Qinghua Jiang

**Author notes:** These authors have contributed equally to this work (Wenyi Yang, Zhaochun Xu, Meng Luo).

## Abstract

With the rapid development of high throughput single-cell RNA sequencing (scRNA-seq) technologies, it is of high importance to identify Cell-cell interactions (CCIs) from the ever-increasing scRNA-seq data. However, limited by the algorithmic constraints, current computational methods based on statistical strategies ignore some key latent information contained in scRNA-seq data with high sparsity and heterogeneity. To address the issue, here, we developed a deep learning framework named DeepCCI to identify meaningful CCIs from scRNA-seq data. Applications of DeepCCI to a wide range of publicly available datasets from diverse technologies and platforms demonstrate its ability to predict significant CCIs accurately and effectively.

## Background

Multicellular life relies on the conformity of cellular activities, which depend on cell-cell interactions (CCIs) across diverse cell types^1–3^. Single-cell RNA-sequencing (scRNA-seq) technologies have enabled remarkable progress in understanding cellular mechanisms at an unprecedented resolution level^4, 5^. Although scRNA-seq data inherently contains gene expression information that could be used to identify intercellular communications, it remains a great challenge to explore potential CCIs that often drive heterogeneity and cell state transitions^6–8^. The signaling events behind cells are usually mediated by interactions of various types of proteins, encompassing ligand-receptor (L-R), receptor-receptor and extracellular matrix-receptor interactions. Especially, multi-subunit L-R complexes are critical for CCIs^9^. Some proteins, such as TGF-beta (transforming growth factor-beta) receptors^10^ and cytokine receptors^11^, require multi-subunit assembly for function^7^. Specifically, in the TGF-beta signaling pathway, the interaction between soluble ligand TGFB1 and the heteromeric complexes of type I and type II receptors (TGFBR1 and TGFBR2) plays an important role in the development of diabetic nephropathy^12^.

To identify CCIs from scRNA-seq data, several computational strategies have been developed based on the L-R gene pairs, such as SingleCellSignalR^13^, iTALK^14^, CellPhoneDB^9^, and CellChat^6^. Each strategy consists of a resource of intercellular interactions prior knowledge and a method to identify underlying CCI events. However, the identified results of these strategies are usually limited by the comprehensiveness of the prior L-R gene pair database. Different L-R pair databases used in each method could contribute to the variety of identified interactions. Also, identifying previously uncharacterized cell types in heterogeneous scRNA-seq data is the precondition for identifying CCIs^15, 16^. Nevertheless, these methods cannot classify cells into cell clusters independently before interaction identification. Moreover, due to the technical difficulties of capturing single-cell proteomic information at present, defining the ground truth of CCI networks is challenging. Recently, deep learning-based methods have demonstrated their prowess in a broad range of single-cell studies. However, there is still no deep learning framework for CCI prediction from scRNA-seq data. Combining scRNA-seq data with deep learning technologies will be greatly expanded to provide unique insights into CCI prediction.

Here, we develop DeepCCI, a graph convolutional network (GCN)^17^-based deep learning framework (https://github.com/JiangBioLab/DeepCCI) for CCI identification from scRNA-seq data. To explore the interactions between cells from scRNA-seq data in one-stop, DeepCCI provides two deep learning models: (i) a GCN-based unsupervised model for cell clustering, and (ii) a GCN-based supervised model for CCI identification. DeepCCI has great potential to cluster cells and capturing biological meaningful interactions between cell clusters by utilizing the underlying complex gene expression patterns of heterogeneous cells from scRNA-Seq data. DeepCCI first learns an embedding function that jointly projects cells into a shared embedding space using Autoencoder (AE)^18^ and GCN. By using the embedding information, DeepCCI clusters cells into several groups. And then, we manually curated a comprehensive signaling-molecule interaction database termed LRIDB for L-R interactions with multi-subunits^9^. According to LRIDB, DeepCCI predicts intercellular crosstalk between any pair of clusters within a given scRNA-seq data. Also, DeepCCI provides several visualization outputs to show how strongly or specifically each cell cluster interacts with every other cell cluster. We demonstrate the overall capabilities of DeepCCI by applying it to several publicly available scRNA-seq datasets. The results indicated that DeepCCI has excellent potential in capturing biological relationships between cells in terms of cell-type clustering and CCI prediction from scRNA-seq data.

## Results

### Overview of the DeepCCI workflow

Given scRNA-seq data, DeepCCI seeks influential representations of cells helpful in performing different tasks in scRNA-seq data analyses (Figure 1). DeepCCI consists of two major functions: (a) clustering cells from scRNA-seq data, and (b) establishing interaction networks between cell clusters with prior knowledge of signaling L-R pairs.

**Figure 1.**
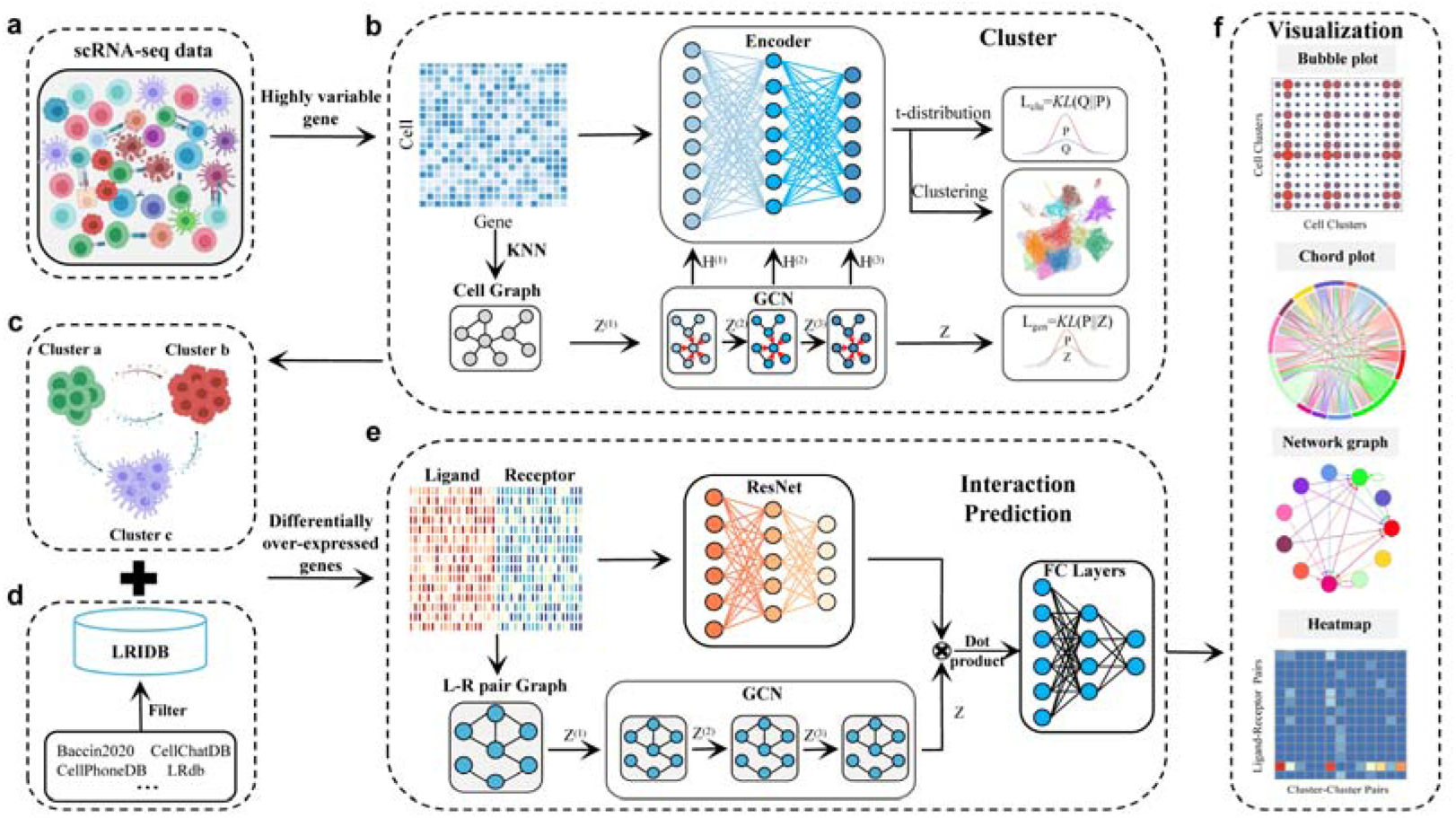
Workflow of the DeepCCI. **a**, DeepCCI takes the scRNA-seq data as input. **b**, DeepCCI clusters cells using the AE and the GCN jointly. **c**, ScRNA-seq data with cell types. **d**, LRIDB contains validated L-R interactions that were collected from several publicly literature-supported databases. **e**, DeepCCI predicts the interactions between cell clusters using ResNet and GCN jointly. **f**, DeepCCI offers several visualization outputs for different analytical tasks.

Cell clustering is a pre-requisite for CCI identification, different number of cell clusters may naturally affect the identified interactions. To establish CCI networks, DeepCCI can operate in label-based and label-free modes. In its label-based mode, DeepCCI requires user-assigned cell labels as the input. In its label-free mode, DeepCCI provides an unsupervised clustering model by using AE and GCN jointly to cluster cells (Figure 1b). The cluster model intakes the gene expression matrix after data preprocessing, including low-quality cells and genes removing, normalization, and variable gene ranking^19, 20^. To achieve better cluster performance, we pretrained an AE model using the top 2,000 variable genes^21, 22^. By minimizing a data reconstruction error, the pretrained AE learns a low-dimensional embedding to reconstruct the gene expression matrix. The pretraining step serves as a prior for the parameter space, which is helpful for the subsequent cluster progress^21^. Then, the pretrained AE was combined with the GCN to build the cluster model. The input of GCN is the cell graph, which is generated from the top 2,000 variable gene expression matrix using the KNN algorithm. The nodes of cell graph represent individual cells while the edges represent neighborhood relations among these cells^23, 24^. Next, the cluster model regenerates the whole graph structure and learns a low-dimensional embedding of each cell jointly using AE and GCN. Based on the learned space embedding, the k-means clustering method was employed to cluster cells^25, 26^.

The more comprehensive set of the signaling L-R pairs is of vital importance to prediction of biologically meaningful intercellular communications. We integrated several public databases, including Baccin2020^27^, Cain2020^28^, CellCallDB^29^, CellChatDB, CellPhoneDB, LRdb^30^, connectomeDB2020^31^, CellTalkDB^32^ and iTALK, into a more comprehensive L-R pairs database named LRIDB (Figure 1d). LRIDB retains 5,127 validated molecular interactions for human and 4,623 for mouse (Supplementary Data 1-2). To investigate the significant interaction between cell clusters, we first identified differentially over-expressed ligands and receptors for each cell cluster based on LRIDB. Next, we associated each interaction with a probability value to quantify interactions between two cell clusters. Specifically, we calculated the interaction probability based on the average expression values of a ligand and the cognate receptor. Three publicly available statistical methods were used to identify the significant interactions (p-value<0.01 or LRscore>0.5). We set these significant interactions as positive samples and the remaining interactions with probability values as negative samples for model training (see “Methods” section). Subsequently, we built a deep learning model for accurately predicting significant CCIs jointly using the Residual Network (ResNet)^33^ and GCN (Figure 1e). The GCN model of this part receives the L-R pair graph that was constructed by the progressed scRNA-seq data and LRIDB. The nodes of L-R pair graph are the L-R pairs, with the edges representing the relationships between these pairs. Simultaneously, the input of ResNet is the expression values of ligands in one cell cluster and those of receptors in another cell cluster. The dot product of outputs from ResNet and GCN was fed into the next fully connected layers^34^ and the significant interactions between cell clusters were predicted (see “Methods” section). In addition, several informative and intuitive visualization methods were used to display the significant connections between interacting cell clusters (Figure 1f).

### Performance evaluation of cluster model of DeepCCI

To evaluate the cell clustering performance of DeepCCI, we compared the cell cluster model of DeepCCI with several state-of-the-art methods (scTAG^35^, Graph-sc^36^, scGNN^37^, scGAE^38^, GraphSCC^39^, scziDesk^40^, scDCC^41^, DCA^18^, DEC^42^, K-means^43^, Spectral) on 12 real-world scRNA-seq datasets (the detailed information is described in Supplementary Data 3) from several representative sequencing platforms. The widely-used clustering metrics Adjusted Rand Index (ARI)^44^ was employed to measure the clustering performance of cluster model of DeepCCI and the other 11 baseline methods (Figure 2a, Supplementary Data 4). To avoid the error caused by chance, each clustering method was run ten times to take the average. For the 12 scRNA-seq datasets, the cluster model of DeepCCI achieved the best ARI on all of them, and even in ‘Qx Limb Muscle’, the ARI reached 0.9848 (Figure 2a, Supplementary Data 4). Meanwhile, Normalized Mutual Information (NMI)^45^ was also used to measure the clustering performance (Fig. S1, Supplementary Data 4). Moreover, 4 major criteria (NMI, Silhouette score^46^, ARI, AMI^47^) were used to evaluate the performance of these ten repeated experiments of DeepCCI cell cluster model (Figure 2b and Fig. S2). The results of these repeated experiments demonstrated the stability of DeepCCI in cell clustering for scRNA-seq data. In addition, we can visualize cell clustering results based on DeepCCI cluster embedding by using the UMAPs^48^ (Figure 2c). By visualizing cell clustering results on UMAPs, one can observe the apparent nearness of cells within the same cluster and separation among different clusters when using embedding provided by the cluster model of DeepCCI (Figure 2c). Our results indicate that the cluster model of DeepCCI can capture the essential hidden information of cells and make full use of the topological relationships among cells to accurately predict cell clusters.

**Figure 2.**
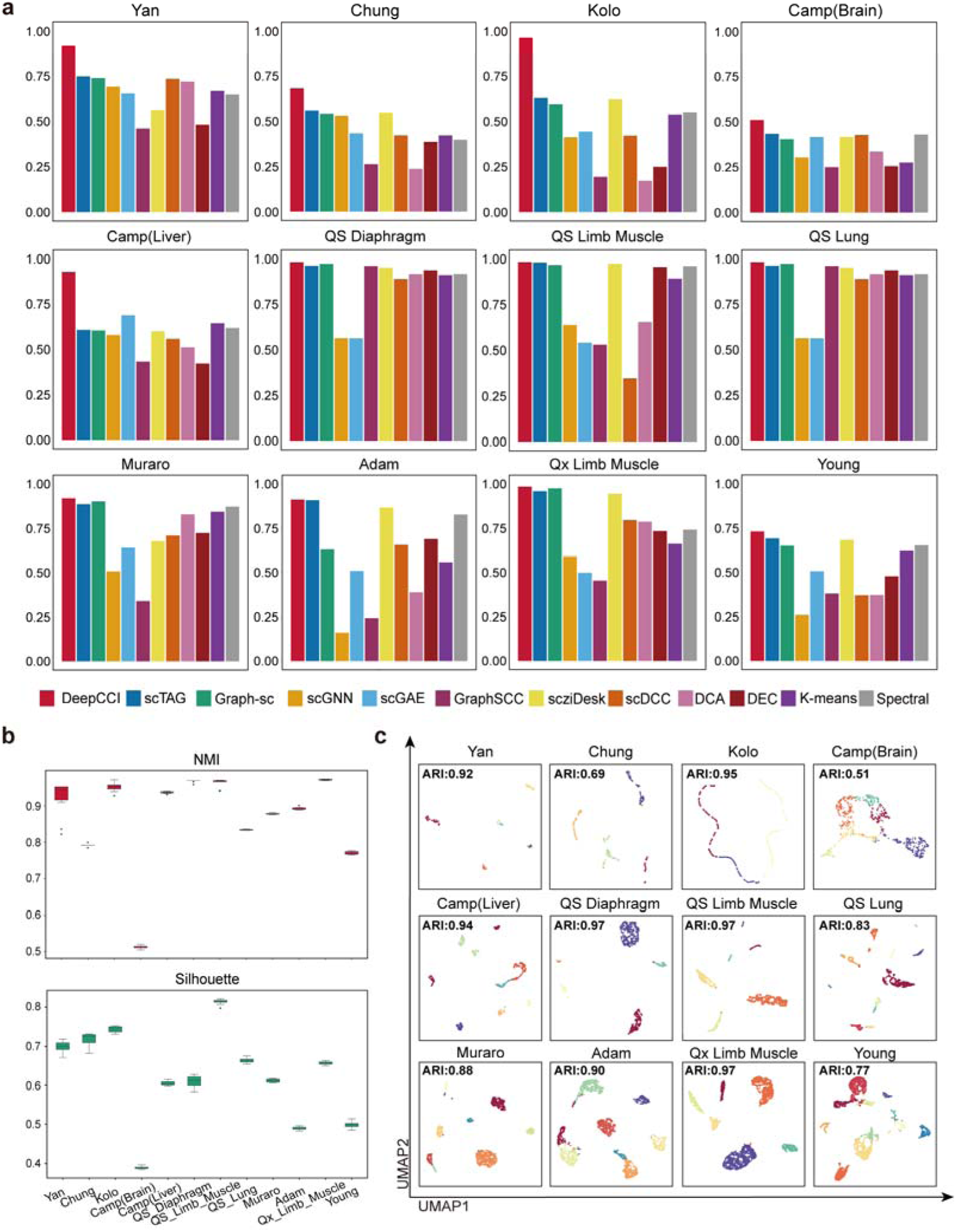
Performance evaluation of cell cluster model of DeepCCI. **a**, Comparison of ARI among cell cluster model of DeepCCI and 11 state-of-the-art methods using 12 real-world scRNA-seq datasets. **b,** Performance evaluations of the cell clustering by repeating cell cluster model of DeepCCI 10 times. **c**, UMAP visualizations for 12 scRNA-seq datasets using the cell embedding generated by cell cluster model of DeepCCI.

On top of that, the cluster number is an important parameter for the performance of unsupervised clustering methods. For scRNA-seq data with unknown cell clusters, the cell cluster model of DeepCCI provides a strategy to determine the number of clusters before clustering and the number of clusters is determined by the Louvain algorithm on the cell graph^25^. To evaluate the effectiveness of this strategy, we continued to compare the clustering performance of DeepCCI cluster model with the several tools (scGNN^37^, Seurat^49^, CIDR^50^, RaceID^51^) that with the ability to automatically define the number of clusters on four common datasets (i.e., Chung^52^, Kolodziejczy^53^, Klein^53^, and Zeisel^54^). In comparison with these methods, DeepCCI has achieved better cluster performance on the overwhelming majority of datasets (Fig. S3). Especially on the Zeisel dataset, the ARI value of the cluster model of DeepCCI is 0.16 higher than scGNN. Taken together, the GCN we used in DeepCCI cluster model can capture the high-order representations of relationships between cells in scRNA-seq data in the context of graph topology, and apply it to the clustering of scRNA-seq data.

### Performance evaluation of interaction model of DeepCCI

To construct an interaction identification model with strong generalization, a published integrated pancreatic islets scRNA-seq dataset (panc8)^49^ across multiple platforms was used. The labeled panc8 data contains eight pancreas datasets across five technologies (CEL-Seq^55^, CEL-Se2^56^, Fluidigm C1^57^, SmartSeq2^58^, inDrops^59^) integrated and normalized with the R toolkit Seurat^60^. The panc8 dataset includes 14,892 cells with 13 benchmark cell labels (Acinar, activated stellate(aPSC), Alpha, Beta, Delta, Ductal, Endothelial, Epsilon, Gamma, Macrophage, Mast, quiescent stellate(aPSC), Schwann) (Figure 3a). To capture the high-confident interactions between cell clusters, we use the benchmark cell labels of the panc8 dataset for further analysis. Based on the LRIDB, we calculate the interaction probability of L-R pairs for the cell cluster pairs and define the significant L-R pairs by using three statistical methods (see “Methods” section).

**Figure 3.**
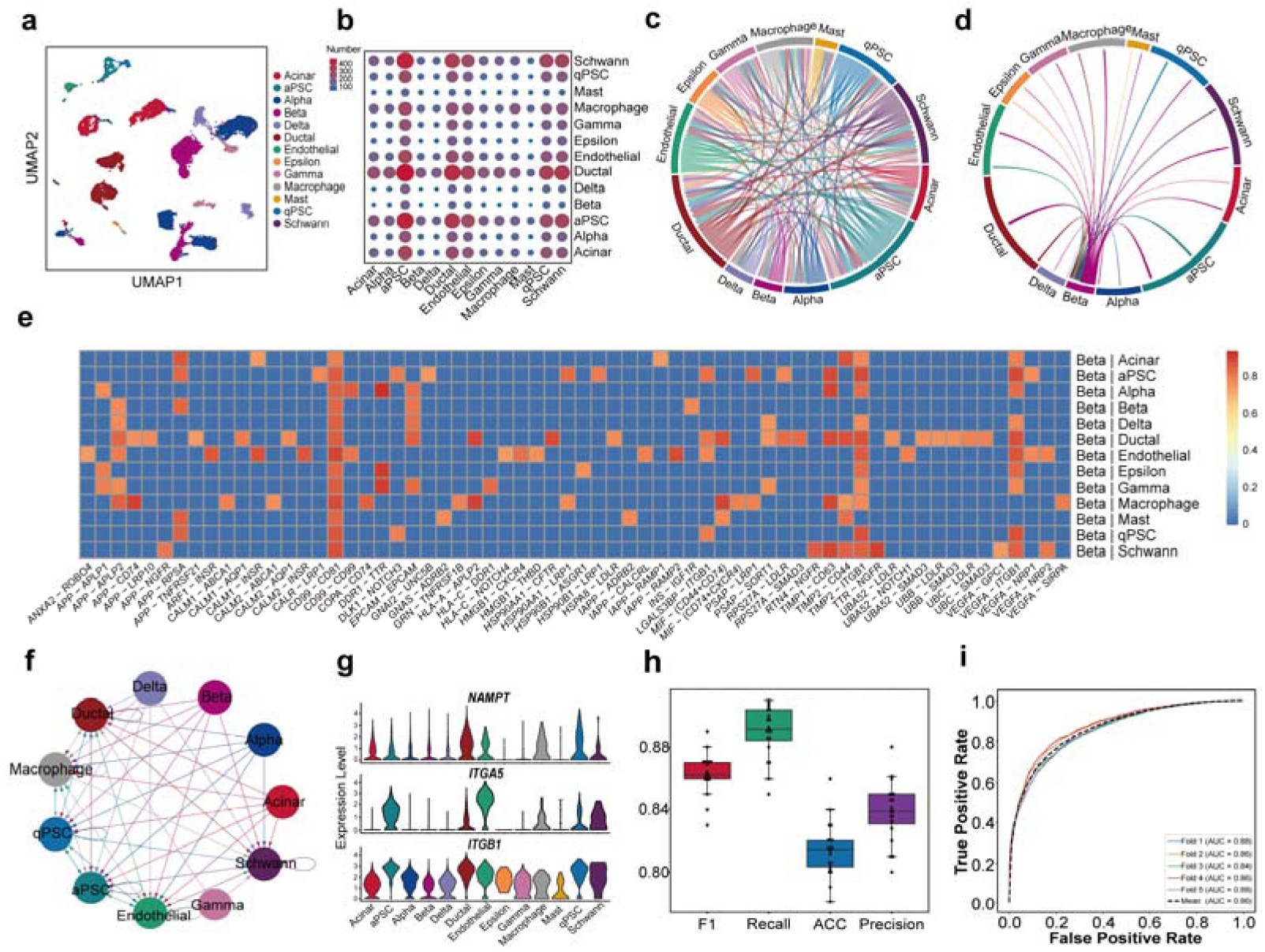
Performance evaluation of cell interaction model of DeepCCI. **a**, Benchmarked cell clusters visualized by UMAP for pancreatic islets scRNA-seq dataset. **b**, Number of significant L-R pairs between any pair of two cell clusters. **c**, Chord plot shows the interactions between all cell clusters. The arrow from source cell cluster point to the target cell cluster and the color of arrow is same as the source cell cluster. **d**, Chord plot shows the interactions from beta cell cluster to other cell clusters. **e**, Heatmap shows the top 200 expressed L-R pairs between beta cluster and other clusters. The color of the heatmap represents the interaction probability value. **f,** Interaction network mediated by the *NAMPT* - (*ITGA5* + *ITGB1*). **g,** Gene expression distribution of the *NAMPT* ligand and *ITGA5*, *ITGB1* receptors. **h**, Performance evaluations of interaction model of DeepCCI of 5-fold cross valuation for 20 times. **i**, The ROC curve of 5-fold cross valuation for DeepCCI interaction model.

To present the significant interactions intuitively, we provide several visualization methods (Figure 3b-g). The bubble plot provides an overview of the number of L-R interactions between cell clusters (Figure 3b). Also, to show all L-R interactions between cell clusters in another intuitive way, we provide the chord plot (Figure 3c-d). The arrows from the source cell cluster in chord plot mean that the ligands in the source cluster are connected with the receptors in the target cell clusters. Not only display the interactions for all cell clusters, chord plot also could show the interactions from one specific cell cluster to other clusters. Different colors in the chord plot represent different cell clusters. Fig 3d shows the interactions between Beta cell and other cell clusters. Paracrine communication between Beta cells and non-Beta cells is known to regulate insulin secretion^61^. The Beta cell is electrically excitable and uses changes in membrane potential to couple variations in the blood glucose concentration to stimulate or inhibition of insulin secretion^62^. And then, the heatmaps were used to reveal the detailed L-R pairs with top 200 probability values from the source cluster to target clusters (Figure 3e). Ligand *MIF* (macrophage migration inhibitory factor) and its receptor *CD74* were found to act as major signaling from Beta cell to Macrophage in pancreatic islet in Figure 3e. Previous study^63^ has claimed that the interaction of *MIF* with the extracellular domain of *CD74* on the cell surfaces can induce activation of the extracellular signal-regulated kinase-1/2 MAP kinase cascade, cell proliferation, and PGE2 production. Meanwhile, additional studies proved that the source of *MIF* seems to be originating from multiple cell types within the islets of Langerhans including, insulin+ beta cells and F4/80+ macrophages^64^. Cells expressing the *CD74* are present on F4/80+ cells within immune cell infiltrate of the islets, and F4/80+ macrophages are the main immune cells in the pancreas expressing *CD74*. These pancreatic macrophages are found in close proximity to insulin+ Beta cells^65^. Through cell-cell contact, Beta cells can direct interaction with the macrophages^66^. Taken together, the defined significant interaction from Beta cells to macrophages mediated by the *MIF*-(*CD74+CD44)* pair and *MIF*-(*CD74+CXCR4)* is in agreement with the reported experimental findings. Moreover, for the L-R pair of interest, the network graph provides an overview of intercellular communication networks, consisting of two components: the cell clusters in the panc8 dataset and the interactions between these clusters (Figure 3f). The color of the edges is the same as the source cluster. The interaction network for the *NAMPT* - (*ITGA5* + *ITGB1*) pair^49^ is shown using a network graph. *NAMPT* - (*ITGA5* + *ITGB1*) pair is from Visfatin signaling pathways^6^ and visfatin was described to be a highly expressed protein with immune cell signaling and nicotinamide adenine dinucleotide (NAD) biosynthetic activity, which is essential for pancreatic beta cell function. Although *ITGB1* subunits are expressed in all cell clusters, the cluster that does not express *ITGA5* subunits is not the target cluster for the source clusters (Figure 3g). Taken together, the standards we used to define significant interactions can capture biologically meaningful CCIs from scRNA-seq data.

Further, based on these significant interactions, the interaction model of DeepCCI was constructed (see “Methods” section). To ensure the generalizability of the interaction model of DeepCCI, we train the model using 5-fold cross-validation. Moreover, 5-fold cross-validation was performed 20 times and five criteria (F1, Recall, ACC, Precision, AUC) were used to measure the performance of interaction model of DeepCCI (Figure 3h-i). Collectively, DeepCCI can identify key features of CCIs within a given scRNA-seq dataset and predict complex intercellular communications in an easily interpretable way.

To prove that the predicted CCI is not biased, DeepCCI was applied to three pancreas datasets^67^ (panc1, panc2, panc3) from different humans obtained using inDrops technologies. These three pancreas dataset includes 1937, 1724 and 3605 cells, respectively. All of these datasets contain 13 benchmark cell labels (Acinar, activated stellate(aPSC), Alpha, Beta, Delta, Ductal, Endothelial, Epsilon, Gamma, Macrophage, Mast, quiescent stellate(aPSC), Schwann)(Fig. S5a-c). The predicted L-R interactions between cell clusters are shown by using chord plot(Fig. S5d-f). Although the three datasets are all derived from the human pancreas, there are significant differences in the predicted CCIs from different datasets. Meanwhile, the interaction network for the *NAMPT* - (*ITGA5* + *ITGB1*) pair^49^ is shown using a network graph (Fig. S5g-i). There are clear differences in the predicted CCIs mediated by *NAMPT* - (*ITGA5* + *ITGB1*) pair from different datasets, some similarities can still be found among them. For example, Endothelial, aPSC and Ductal are the main target cell types mediated by *NAMPT* - (*ITGA5* + *ITGB1*) pair. In addition, the detailed L-R pairs with top 200 probability values from Beta cells to other cell types are shown using the heatmap (Fig. S5j-l). Due to the differences that exist between samples, the predicted CCIs between Beta cells and other cell clusters from the three datasets show different results. Meanwhile, the predicted interactions between Beta cells and Ductal cells from panc1 dataset and panc3 dataset have a large similarity. This is in agreement with a reported experimental finding^67^. Taken together, DeepCCI has strong generalization and robustness and can fully utilize the information existing in the scRNA-seq data.

### Independent testing of interaction model of DeepCCI

To investigate whether the interaction model of DeepCCI is applicable to the prediction of CCIs using scRNA-seq data, we used two different datasets (10X Genomics): human atopic dermatitis (AD) dataset^67^ and embryonic mouse skin dataset^68^. The human AD dataset contains 17349 cells which were clustered into 12 cell groups, including T cells (TC), APOE + fibroblast (FIB), CD40LG + TC, COL11A1 + FIB, FBN1 + FIB, inflammatory dendritic cells (Inflam.DC), Inflam.FIB, Inflam.TC, Langerhans cells (LC), Natural Killer T cells (NKT), cDC1, cDC2. The cell clusters for the two datasets are visualized using the UMAP method (Figure 4a). The embryonic mouse skin dataset contains 25148 cells, which were clustered into 13 cell groups, including Basal, Basal-P, DC, endothelial (ENDO), FIB-A, FIB-B, FIB-P, Immune, melanocyte (MELA), myeloid (MYL), Muscle, Pericyte, Spinious (Figure 4b). To verify the generalizability of DeepCCI interaction model independently, we used the benchmarked cell labels of these two datasets to predict significant CCIs. Four criteria are used to evaluate the performance of the model (Figure 4c). These results confirm the generalizability of DeepCCI in the prediction of CCIs on diverse scRNA-seq datasets. Also, several visualization methods are used to show the predicted interaction diversification (Figure 4c-j).

**Figure 4.**
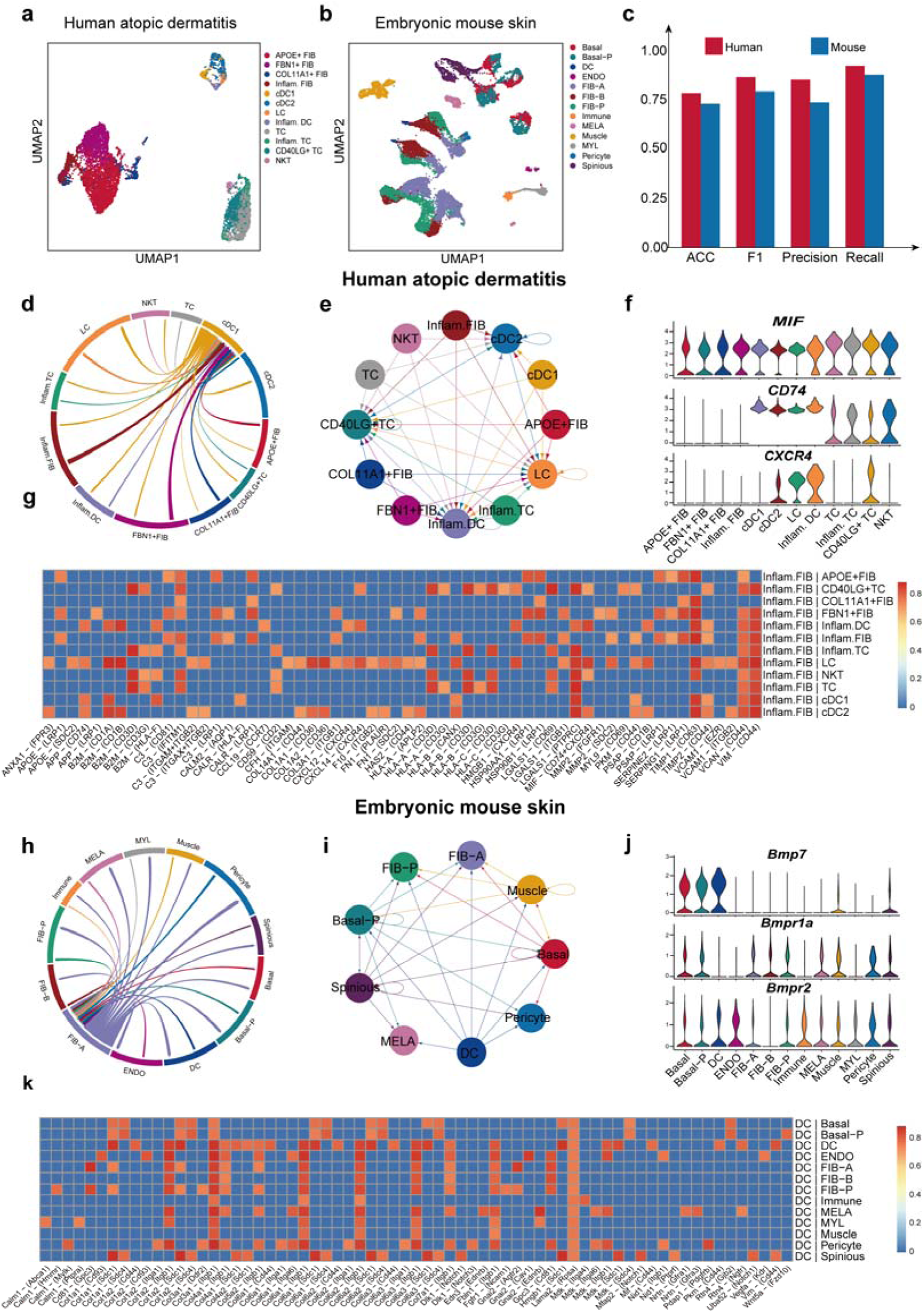
Independent testing of interaction model of DeepCCI. **a-b**, Benchmarked cell clusters visualized by UMAP. **a** represents the human atopic dermatitis dataset and **b** represents embryonic mouse skin dataset. **c**, Performance of interaction model of DeepCCI on human and mouse scRNA-seq datasets. **d,** Chord plot shows the predicted interactions from cDC1 to other cell clusters for the human atopic dermatitis dataset. **e**, The predicted interaction network mediated by *MIF* - (*CD74* + *CXCR4*). **f**, Gene expression distribution of the *MIF*, *CD74* and *CXCR4*. **g**, Heatmap shows the top 200 expressed predicted L-R pairs for Inflam.FIB to other clusters. **h,** Chord plot shows the predicted interactions from FIB-A cell cluster to other clusters for the embryonic mouse skin dataset. **i**, Interaction network mediated by *Bmp7* - (*Bmpr1a* + *Bmpr2*) pair. **j**, Gene expression distribution of the *Bmp7*, *Bmpr1a* and *Bmpr2*. **k**, Heatmap shows the top 200 expressed predicted L-R pairs for DC to other clusters.

Atopic dermatitis (AD) is a prevalent inflammatory skin disease with a complex pathogenesis, involving immune cell and epidermal abnormalities^69^. For the human AD dataset, the number of predicted L-R pairs between the cell clusters for these different datasets is shown using the bubble plot (Fig. S6). The chord plot shows the interactions between cDC1 cell cluster and other clusters for the human AD dataset (Figure 4d). cDC1 and cDC2 were divided from the dermal conventional dendritic cells. Characterized by expression of XCR1 and cross-presentation to CD8+ T cells, cDC1s are involved in immunity to intracellular pathogens and cancer immunity^70^. FIBs are found in most tissues, yet they remain poorly characterized. Different fibroblast subpopulations with distinct functions have been identified in the skin. In addition to autocrine signals, FIBs are highly responsive to Wnt-regulated signals from the overlying epidermis, which can act both locally, via extracellular matrix (ECM) deposition, and via secreted factors that impact the behavior of fibroblasts in different dermal locations^71^. Meanwhile, the intercellular network of the *MIF* - (*CD74* + *CXCR4*) pair for the human disease skin dataset (Figure 4e-f). MIF has been implicated in the progression of many inflammatory and autoimmune diseases^72–74^. MIF activities are mediated by non-cognate interactions with the *CXC* chemokine receptors *CXCR2* and *CXCR4* or by ligation of *CD74*, which is the cell surface-expressed form of class II invariant chain^75^. Previous studies identified the MHC class II chaperone *CD74*, the cell surface form of class II invariant chain as well as the chemokine receptors *CXCR2* and *CXCR4* as receptors for *MIF* on different cell types. Moreover, the top 200 probability values of predicted L-R pairs from Inflam.FIB cluster to other cell clusters for the human AD dataset (Figure 4g). DeepCCI predicted L-R pair *CCL19*-*CCR7* as the significant signaling for the communication from Inflam.FIB to Inflam.DC (Figure 4g). This is in agreement with a reported experimental finding^76, 77^. In addition, FIBs are a heterogeneous population of cells that play a critical role in the dynamics of ECM composition and architecture^78, 79^ and *CD44* is a cell surface adhesion receptor expressed on nearly all cell types present in the dermis. The previous study has proved that *CD44* was required for directional migration of fibroblasts in an in vitro wound healing model. Collagen is a *CD44* ligand^80^ generated after cleavage from the extracellular membrane, can be integrated and bind to ECM components including collagen^81^. *CD44* plays an important role in fibrillar collagen accumulation and wound healing during the injury response through interacting with collagen ligands. Therefore, the interactions predicted by DeepCCI between type I collagen ligand (*COL1A1* and *COL1A2*) and *CD44* in the human AD dataset have biological significance. In addition, LCs are migratory cells and continually travel to the skin draining lymph nodes to promote tolerance in homeostasis^82^ or to initiate adaptive immune responses^83^. The migration of LCs through the dermis is mediated via *CXCR4* signaling after binding to its cognate chemokine *CXCL12* produced by FIBs and the interaction has been predicted by DeepCCI.

For the embryonic mouse skin dataset, the interactions between FIB-A cell cluster and other clusters by using chord plot (Figure 4h). Simultaneously, *Bmp7* - (*Bmpr1a* + *Bmpr2*) pair are displayed by the network graphs (Figure 4i-j). The *Bmp7*-(*Bmpr1a*+*Bmpr2*) pair is from Bone Morphogenetic Protein (BMP) pathway. The BMP pathway is a critical regulator of development and belongs to the cytokine growth factor TGF-beta family^84^. BMP activity has largely been viewed as tumor-suppressive as demonstrated by the loss and gain of function of BMP signaling components. The TGF-beta family ligand *Bmp7* shows a tightly regulated inverse expression patterns within the epidermal microenvironment during steady-state and inflammation^85^. When *Bmpr2* is expressed as a dominant-negative in a mouse model of breast cancer, it enhances tumor metastasis through a paracrine inflammatory microenvironment^86^. The top 200 probability values of predicted L-R pairs from the DC cluster to other cell clusters for the embryonic mouse skin dataset are shown using the heatmap (Figure 4k). Dendritic cells play a very important role in activating both the innate and adaptive immune systems upon acute injury to tissue. DCs serve as messengers between the innate immune response and the adaptive immune response, a process that is active in each phase of wound healing. Our predicted L-R interactions showed strong intercellular communication between the DC cluster and other cell clusters including the Notch signaling ligands *Dlk1*, and collagens family members *Col1a1, Col1a2, Col4a1* (Figure 4k). The overexpressed Collagen ligands of DCs and their interaction with receptors from other cell types uncover the important role of DCs in sensing tissue damage and promoting wound repair in the skin. Taken together, for datasets generated by different platforms, DeepCCI has the ability to accurately identify meaningful interactions from complex and highly sparse scRNA-seq data.

### Comparison of DeepCCI with state-of-the-art methods for predicting CCIs

We systematically compared DeepCCI with seven state-of-the-art methods (SingleCellSignalR, iTALK, CellPhoneDB, CellChat, CellCall, CytoTalk and NATMI) that offered thresholds for communication scores and all the methods were applied to infer cell-cell interactions using their own default parameters. To demonstrate the performance of DeepCCI from multiple aspects, we use three datasets from different techniques (a scRNA-seq dataset of human testicular^87^, a seqFISH dataset of mouse organogenesis^88^ and 10X Visium spatial transcriptomics dataset of coronal section of the mouse brain^89^).

First, we compared the performance of DeepCCI with seven CCI prediction methods on the dataset of human testicular cells. The curated literature related CCIs from Sertoli cells (STs) to spermatogonial stem cells (SSCs) are provided as the ground truth^29^. DeepCCI identified 47 CCIs from STs to SSCs and SingleCellSignalR, iTALK, CellPhoneDB, CellChat, CellCall, CytoTalk and NATMI identified 93, 22, 54, 49, 47, 30 and 107 CCIs by their own default cut-offs (Fig 5a, see Supplementary Data 6 for details). Over 89% of DeepCCI-identified CCIs (42/47) have been reported to be involved in spermatogenesis and the high literature support rate is superior to other tools. The area under curve of receiver operating characteristic (AUC) and precision was used to compare these methods (Figure 5b). DeepCCI achieved the highest AUC and precision on the scRNA-seq dataset of human testicular. These results suggest that DeepCCI has the ability to discover biologically meaningful CCIs from scRNA-seq data.

**Figure 5.**
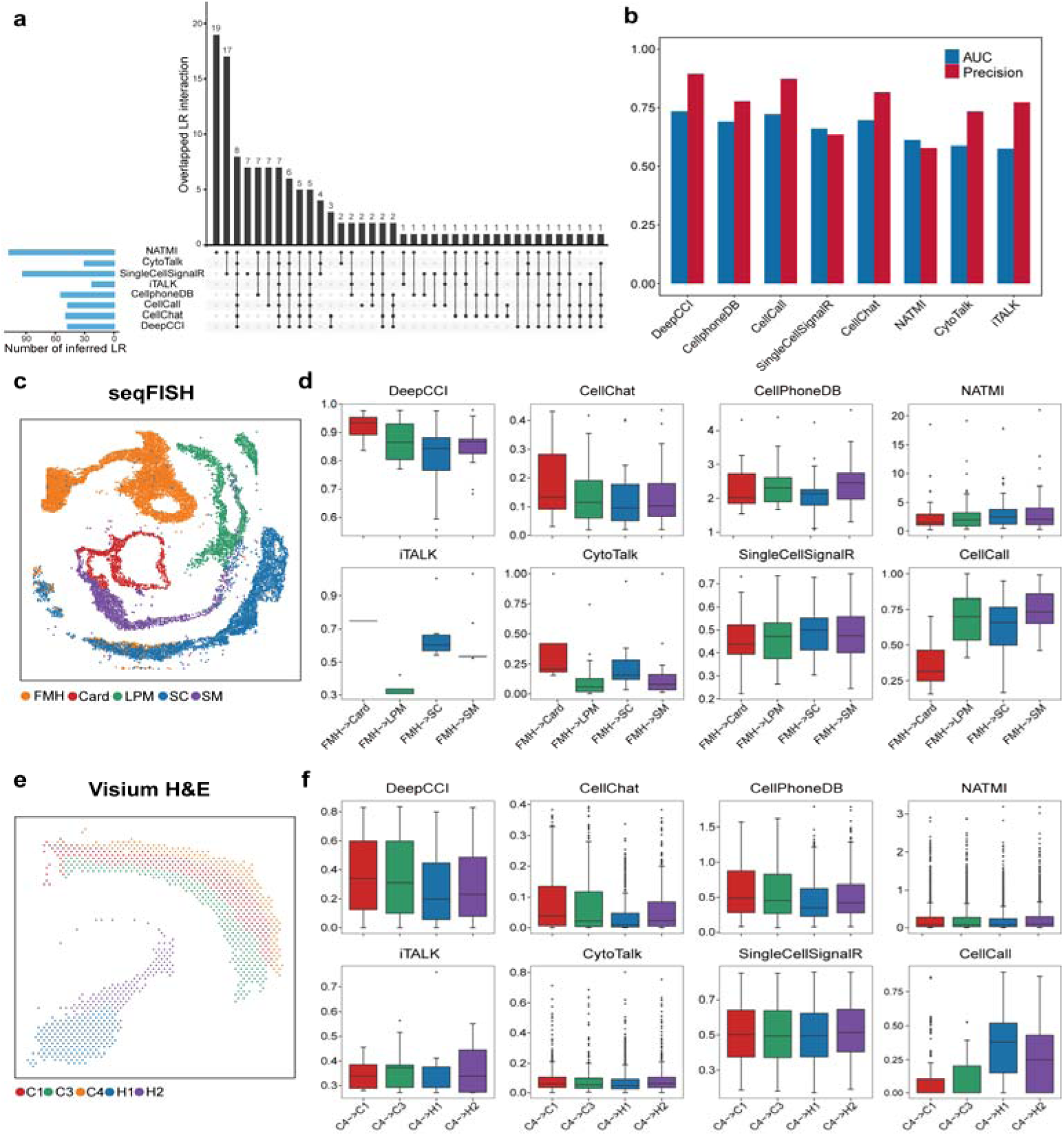
Comparison of the performance of DeepCCI with state-of-the-art methods. **a,** UpSetR plot of predicted CCIs from the eight tools of human testicular cells. **b,** The AUC and precision of eight CCI predicted methods. **c**, Benchmarked cell clusters visualized by UMAP for the seqFISH dataset of mouse organogenesis. **d,** Comparison of the interaction strength between spatially adjacent and distant cells for the seqFISH dataset. The interaction strength is calculated by the interaction probability or score. **e,** Benchmarked cell clusters visualized by UMAP for 10X Visium spatial transcriptomics dataset of the mouse brain. **f,** Comparison of the interaction strength between spatially adjacent and distant cells for 10X Visium spatial transcriptomics dataset of the mouse brain.

Next, we leveraged spatial information as a way to support the predictive potential of DeepCCI, under the assumption that spatially adjacent cell types are expected to have stronger cell-cell interaction than other non-adjacent cell types. For the seqFISH dataset of mouse organogenesis, we made use of the single-cell resolutions of seqFISH dataset to identify both the spatially adjacent cell types and spatially distant cell types to obtain the interaction predictions. To more intuitively show the interaction differences between cell types at different distances, five cell types with obvious clustering were selected in this case (Figure 5c), including Lateral plate mesoderm (LPM), Forebrain/Midbrain/Hindbrain (FMH), Spinal cord (SC), Splanchnic mesoderm (SM), Cardiomyocytes (Card). Then, we selected the FMH as the source cell cluster and computed the interaction probabilities or scores between FMH and other cell clusters (Figure 5d). According to the spatial location of these cells, the Crad cells are spatially adjacent to FMH and the SC cells are spatially distant to FMH. Also, LPM cells are slightly closer to FMH in spatial position than SC cell type. We compared the interaction strength (probabilities or scores) between each cell cluster predicted by DeepCCI with the other seven methods (Figure 5d). We found that DeepCCI consistently captures stronger interactions in spatially adjacent cells than in distant cells of the interaction probabilities. Meanwhile, DeepCCI also performed well at discriminating spatially adjacent from distant cells and other tools failed to capture stronger interactions in spatially adjacent cells as compared to spatially distant cells.

We conducted a similar analysis with 10X Visium spatial transcriptomics dataset of the mouse brain. In this case, five cell types were selected to study the interactions between cell clusters (Figure 5e), including Cortex_1 (C1), Cortex_3 (C3), Cortex_4 (C4), Hypothalamus_1 (H1), Hypothalamus_2 (H2). The C4 cell type was selected as the source cell type and the interaction probabilities or scores from C4 cell type to other cell types were computed. From the spatial distribution of these cell types, we can easily find the spatial relationship of C4 cell type with other cell types. The C1 and C3 cell types are spatially adjacent to C4 cell type and the H1 and H2 cell types are spatially distant to C4 cell type. In addition, compared with H1 cell type, H2 is slightly closer to C4 cell type in spatial position (Figure 5e). We found a clear association between the predicted CCIs and the spatial adjacency of their corresponding cell types for DeepCCI, CellChat and CellPhoneDB, while the other methods showed inconsistent trends (Figure 5f). Moreover, compared with CellChat and CellPhoneDB, the predicted interactions of DeepCCI from C4 cell type to C3 cell and H2 cell are better to reflect the spatial relationship between C4 and these two cell types. Together, our analyses show that DeepCCI performs well at predicting biologically meaningful interactions in spatially adjacent cells than in distant cells from spatial transcriptomics datasets.

### De novo prediction of cell-cell interactions using DeepCCI

To test the complete ability of DeepCCI for cell clustering and interaction prediction, we applied DeepCCI on a scRNA-seq dataset called PBMC3k^90^ generated using 10×Genomics technology. PBMC3k contains 2700 cells with 9 benchmarked labels (Figure 6a). Firstly, we clustered the cells to 9 groups using the cluster model of DeepCCI and ARI was employed to measure the clustering performance. The ARI value reached 0.71 and the visualization result is shown using UMAP (Figure 5b). Next, to display the predicted cell clusters more intuitively, the scCATCH^91^ was used to labeled the predicted cell clusters (Figure 6b). From the visualization result, we can see that the annotated labels are roughly the same as the benchmarked labels. Specifically, most Memory CD4 T cells were clustered into two groups by DeepCCI cluster model and annotated to CD1C+_B dendritic cell (CD1C+_B DC) and Circulating Fetal cell by scCATCH. That is because the number of Platelet cells is very small (only 14 cells) and these cells were clustered to the Naive CD4 T cells. By comparing the expression of marker genes, it is easy to find the gene expression of CD1C+_B dendritic cells are almost identical to the Circulating Fetal cells (Fig. S10). It means that the interactions associated with these two cell types will be similar, and these CCIs will be similar as well to Memory CD4 T-related CCIs (Figure 6g-j). Finally, we identified the significant interactions jointly using these predicted cell clusters and the interaction model of DeepCCI.

**Figure 6.**
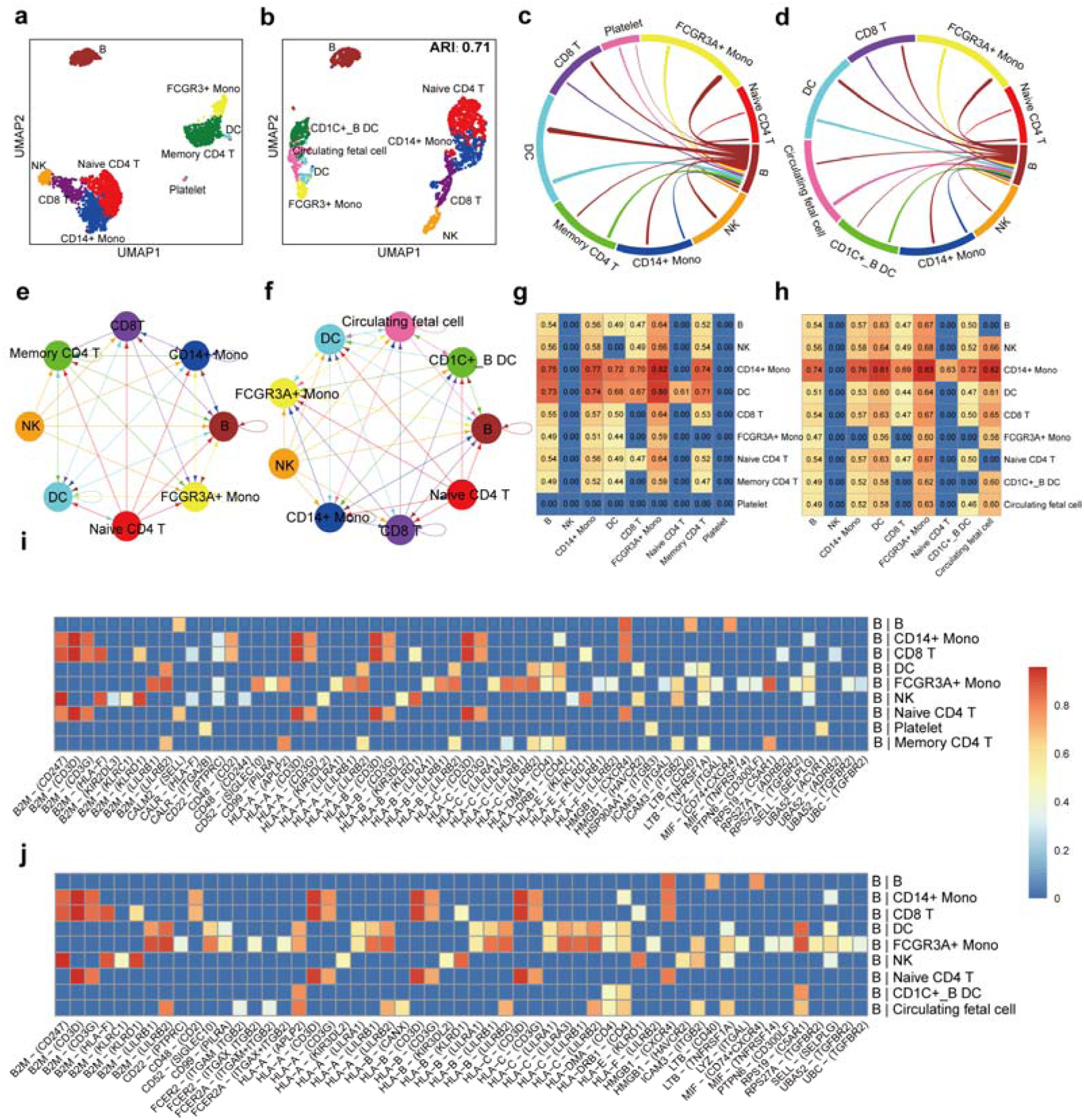
Comparison analysis between high-confidence and predicted interactions. **a,** Benchmarked cell labels of PBMC3K dataset visualized by UMAP. **b,** Predicted cell labels of PBMC3K dataset visualized by UMAP. **c**, High-confidence interactions between B cell cluster and other cell clusters. **d,** Predicted interactions from B cell cluster to other clusters. **e,** High-confidence interaction network mediated by *MIF* - (*CD74* + *CD44*) pair. **f**, High-confidence interaction network mediated by *MIF* - (*CD74* + *CD44*) pair. **g,** Interaction probability values of high-confidence interaction between all clusters under the *MIF* - (*CD74* + *CD44*) pair. **h,** Interaction probability values of predicted interactions between all clusters under the *MIF* - (*CD74* + *CD44*) pair. **i,** Heatmap of high-confidence L-R pairs with the interaction probability value for B cell cluster to other clusters. **j,** Heatmap of predicted L-R pairs with the interaction probability value for B cell cluster to other clusters.

For ease of comparison, we set the interaction results defined by benchmarked labels and three statistical methods (see “Methods” section) as the high-confidence interactions. Then, we compared predicted interactions with the high-confidence interactions. The chord plots display the interactions from the B cell cluster to other cell clusters for the two conditions (Figure 6c). And then, the interactions mediated by *MIF* - (*CD74* + *CD44*) are shown using network graph and the gene expression of L-R pair for all clusters is detailed shown (Figure 6e-f). *MIF* activates the ERK (extracellular signal-regulated kinase) 1/2 MAPK (mitogen-activated protein kinases) signaling pathway by binding to the *CD74*/*CD44* complex, which appears to have an accessory role in immune cell stimulation^92^. The phosphorylation of MAPK, upon engagement of *CD74*, requires the co-expression of *CD44*, a known activator of the Src tyrosine kinase^93, 94^. To show the interactions mediated by *MIF* - (*CD74* + *CD44*) clearly, the interaction probability values between cell clusters were shown (Figure 5g-h). By comparing these two results, we observed that the predicted interaction network mediated by *MIF* - (*CD74* + *CD44*) pair is generally consistent with the high-confidence interactions. Our results confirmed that DeepCCI can accurately predict important CCIs under the condition of similar cell clusters. Moreover, the heatmaps show the interaction probability of L-R pairs from the B cell cluster to other clusters for high-confidence interactions and predicted interactions, respectively (Figure 6i-j). The resulting heatmap of predicted interactions is similar to high-confidence interactions suggesting DeepCCI could better highlight significant interactions. CD48 is a member of the signaling lymphocyte activation molecule family and participates in the adhesion and activation of immune cells^95^. Given the broad expression pattern of CD48, and its increased expression under inflammatory conditions, CD48 is positioned to influence many immunological processes by both cell-intrinsic and ligand-receptor actions. CD48 can have activating roles on T cells, antigen-presenting cells and granulocytes, by binding to CD2 or bacterial FimH, and through cell-intrinsic effects. CD2 is predominantly expressed on T and natural killer (NK) cells in human^96, 97^. CD2 and CD48 enhance early T cell signaling and CD2 was required for CD48 to associate with the TCR and CD3. In addition, a previous study^95^ has proved the interactions of CD48 and CD2 contribute to both priming and effector functions of CD8+ T cells. Therefore, the predicted CD48-CD2 interaction by DeepCCI from B cell to Naïve T cells and CD8+ T cells have biological significance. Moreover, Naïve CD8 T cells become activated when they recognize peptide antigen bound to major histocompatibility complex-I (MHC-I) presented by professional antigen-presenting cells (APCs). Part of the TCR-CD3 complex present on the T-lymphocyte cell surface plays an essential role in the adaptive immune response^98^. When APCs activate the TCR, TCR-mediated signals are transmitted across the cell membrane by the CD3 chains CD3D, CD3E, CD3G and CD3Z (Figure 6j). Beta-2-microglobulin (B2M) is the component of the MHCI and is involved in the presentation of peptide antigens to the immune system^99^. Thus, the predicted interactions between overexpressed CD3 receptor in CD8 T cells and the ligand expressed in B cells mean CD8 T cells are in a state of the immune response. In addition, the interactions of human leukocyte antigen(HLA)-I ligand (HLA-A, HLA-B, HLA-C) from B cells and CD3 receptor also reflects this process (Figure 6j). The above immune response process can obtain theoretical support from the T cell receptor signaling pathway. Compared with high confidence interactions defined by multiple statistical methods, the interaction model of DeepCCI can predict significant interactions both in general and in specific cases. Taken together, DeepCCI has the ability to identify CCIs from scRNA-seq data in one stop. Even for scRNA-seq data without cell labels, DeepCCI can capture biologically meaningful cell clusters and identify the significant interactions between these clusters.

## Discussion and conclusion

It is still a fundamental challenge to identify intercellular communications from high-volume, high-sparsity, and noisy scRNA-seq data. We described DeepCCI, a tool to help users identify cell clusters and CCIs from scRNA-seq data. To cluster cells from scRNA-Seq data, DeepCCI maps all cells in a joint low-dimensional embedding space using GCN-based deep neural networks. The learned joint embedding can be treated as the high-order representations of relationships between cells in scRNA-Seq data. In comparison to several state-of-the-art methods for scRNA-seq clustering, DeepCCI achieved the best clustering accuracy across various datasets. To explore more biologically meaningful intercellular interactions from scRNA-seq data, we constructed the most comprehensive set of L-R pairs. DeepCCI is the first approach to explore intercellular interactions using the deep learning framework from scRNA-seq data. The key innovations of DeepCCI are incorporating global propagated topological features between L-R pairs through GCNs, together with integrating gene expression of cell pairs in the prediction process for intercellular interactions. DeepCCI allows users to quickly identify significant interactions based on single-cell expression data and visualize overall CCI networks under different conditions.

Although we defined the significant CCIs based on the most comprehensive L-R pairs database and several publicly available statistical methods, systematic evaluation of predicted CCIs is challenging due to the lack of ground truth. Recent advances in spatially resolved transcriptomic techniques^100^ offer an opportunity to explore the spatial organization of cells in tissues. Integrating scRNA-seq and spatial transcriptomics data may therefore increase our understanding of the roles of specific cell subpopulations and their interactions in development, homeostasis and disease^101^. In the future, we will continue to enhance DeepCCI to support the integration of single-cell multi-omics sequencing (scMulti-seq) data and incorporate spatial transcriptomics to make the analyses more explainable. We plan to develop a more user-friendly software system based on our DeepCCI model, together with modularized analytical functions in support of data format standardization, quality control, data integration, multi-functional scMulti-seq analysis, performance evaluation, and interactive visualization.

## Methods

### Database construction for ligand-receptor pairs

To construct a database of L-R interactions that comprehensively represents the current state of knowledge, we manually reviewed publicly available L-R interaction databases and developed LRIDB. LRIDB is a database of L-R interactions in both mouse and human. Detailed, we collected L-R interactions from publicly available databases (Baccin2020, Cain2020, CellChatDB, CellCallDB, CellPhoneDB, LRdb, connectomeDB2020, CellTalkDB and iTALK). These databases contain L-R pairs with the literature support for human and mouse. The Baccin2020, Cain2020 contain the L-R interactions for mouse. The CellPhoneDB, LRdb, connectomeDB2020 and iTALK contain the L-R interactions for human. Meanwhile, CellChatDB, CellTalkDB and CellCallDB contain L-R interactions for both human and mouse. We manually select L-R pairs from these databases and only retain the most comprehensive interaction for the shared interactions of these databases.

### Dataset preprocessing

DeepCCI takes the scRNA-Seq gene expression profile as the input. Data filtering and quality control are the first steps of data preprocessing, only genes expressed as non-zero in more than 1% of cells, and cells expressed as non-zero in more than 1% of genes are kept.

For the prediction of cell clusters, we used Seurat to normalize each cell to 10,000 read counts and log-transformed the data. Then, genes are ranked by standard deviation, i.e., the top 2000 genes in variances are used for the study.

For the study of the interactions between cell clusters, we adopted another different data processing method. First, we identified differentially expressed signaling genes across all cell clusters within a given scRNA-seq dataset, using the Wilcoxon rank sum test^102^ with a significance level of 0.05. Next, we project gene expression data onto a high-confidence experimentally validated protein-protein interaction (PPI) network from STRINGdb^103^. Overexpressed L-R pairs were selected if a protein of each interaction was in the list of ortholog ligands and the other was in the list of ortholog receptors. The obtained gene expression data are used for the definition and prediction of interaction between cell clusters. To reduce the effects of the extreme values, we calculated the ensemble average expression of ligands and receptors in a given cell cluster using a truncated mean method and 10% was selected as the truncation ratio.

### Identification of statistically significant interactions

We calculate the interaction probability value of L-R mediated cluster-cluster interactions by using the expression product of two cell clusters. Based on the projected data progressed, the interaction probability P_i,j_ from cell clusters **i** to **j** for a particular L-R pair k was modeled by:

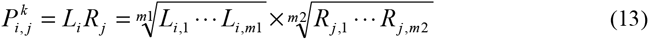

where L_i_ and R_j_ represent the average expression level of ligand L and receptor R in cell group **i** and cell group **j**, respectively. The expression level of ligand L with m1 subunits (i.e., L_i,1_, □, L_i,m1_) was approximated by their geometric mean, implying that the zero expression of any subunit leads to an inactive ligand. Similarly, we computed the expression level of receptor R with m2 subunits.

Then, the significant interactions between two cell clusters are identified using the publicly available statistical methods (CellChat, CellPhoneDB, SingleCellSignalR). Specifically, we applied these methods to the single-cell datasets by using the same L-R interactions database, LRIDB, to identify the statistically significant interactions. By using the threshold (LRscore >= 0.5 or p-value <= 0.01) for each method, we selected the statistically significant interactions as the true labels of the model. Specifically, to explore more meaningful interactions, the majority vote was used to select the significant interactions.

### Autoencoder (AE) and Graph Convolutional Network (GCN)

The basic AE is employed to learn the representations of the single-cell expression data in order to accommodate for different kinds of data characteristics. We assume that there are *L* layers in the AE and *l* represents the layer number. Specifically, the representation learned by the *l*-th layer in encoder part, *H*^(l)^, can be obtained as follows:

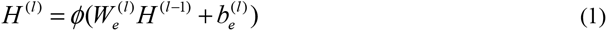

where *ϕ* is the activation function of the fully connected layers, 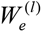 and 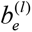 are the weight matrix and bias of the *l*-th layer in the encoder, respectively. Besides, we denote *H*^(0)^ as the pre-progressed single-cell expression data X.

The encoder part is followed by the decoder part, which is to reconstruct the input data through several fully connected layers by the equation:

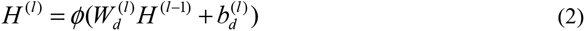

where 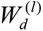 and 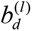 are the weight matrix and bias of the *l*-th layer in the decoder.

The output of the decoder part is the reconstruction of the feature data X, which results in the following objective loss function:

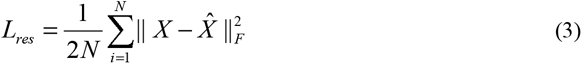

where *X̂* is the reconstructed expression data by the AE.

The GCN is employed to explore the underlying topology information of the cell graph and L-R pair graph. The GCN module is used to accommodate two different kinds of information, i.e., data itself and relationship between data. The essential idea is to update the node representations by propagating information between nodes. With the weight matrix *W*, the representation learned by the *l*-th layer of GCN, *Z*^(*l*)^, can be obtained by the following convolutional operation:

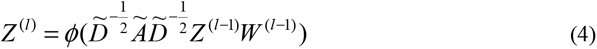

where 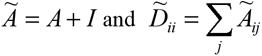. *I* is the identity diagonal matrix of the adjacent matrix *A* for the self-loop in each node. The representation *Z*^(*l*–1)^ will propagate through the normalized adjacency matrix 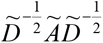 to obtain the new representation *Z*^(*l*)^.

### Model construction for single-cell clusters

To infer the cluster of single cells, we first built the cell graph from a KNN graph, where nodes are individual single cells, and the edges are relationships between cells. K (default = 10) is the predefined parameter used to control the scale of the captured interaction between cells. Each node finds its neighbors within the K shortest distances and creates edges between them and itself.

Considering that the representation learned by AE *H*^(*l*)^ is able to reconstruct the feature itself and contains different valuable information, we combine the two representations *Z*^(*l*)^ and *H*^(*l*)^ together to get a more complete and powerful representation as follows:

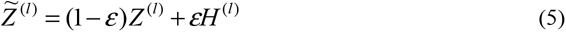

where ε is a coefficient to balance the weights of GCN and AE, and we uniformly set it to 0.5 here. In this way, we connect the AE and GCN layer by layer to get the more informative embedding for the next clustering progress.

Then we use *Z̃*^(*l*)^as the input of the *l*-th layer in GCN to generate the representation *Z̃*^(*l* +1)^:

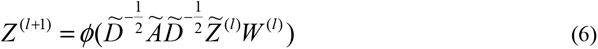

the AE-specific representation *H*^(*l*)^ will be propagated through the normalized adjacency matrix 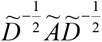. To preserve information as much as possible, we transfer the representations learned from each GCN layer into a corresponding encoder layer and decoder layer for information propagation. Now, we have connected the AE with GCN in the neural network architecture. Here, we unify the AE and GCN modules in a uniform framework and effectively trains the two modules end-to-end for clustering. Learning an effective data representation is of great importance to deep clustering. Pretraining is a required step of the DeepCCI model, and it gives a substantial boost in the performance compared to starting from the random weights.

In particular, for the *i*-th cell and *j*-th cluster, we use the Student’s t-distribution as a kernel to measure the similarity between the data representation *h_i_* and the cluster center vector *µ_j_* as follows:

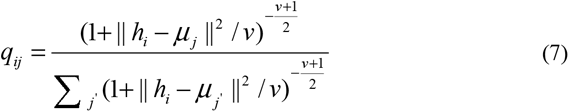

where *h_i_* is the *i*-th row of H^(L)^, *µ_j_* is initialized by K-means on representations learned by pretrained AE and *v* are the degrees of freedom of the Student’s t-distribution. *q_ij_* can be considered as the probability of assigning cell *i* to cluster *j.* We treat Q = [*q_ij_*] as the distribution of the assignments of all samples and let α=1 for all experiments.

After obtaining the clustering result distribution Q, we aim to optimize the data representation by learning from the high confidence assignments. Specifically, we want to make data representation closer to cluster centers, thus improving the cluster cohesion. Hence, we calculate a target distribution P as follows:

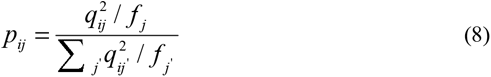

where 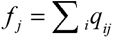 are soft cluster frequencies. In the target distribution P, each assignment in Q is squared and normalized so that the assignments will have higher confidence, leading to the following objective function:

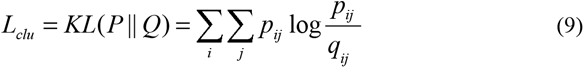

By minimizing the KL divergence loss between Q and P distributions, the target distribution P can help the encoder learn a better representation to make the data representation surrounding the cluster centers closer. The target distribution P is calculated by the distribution Q, and the P distribution supervises the updating of the distribution Q in turn.

The GCN module will also provide a clustering assignment distribution Z. Therefore, we can use distribution P to supervise distribution Z as follows:

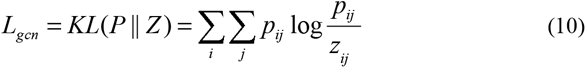

The overall loss function of the cluster model is:

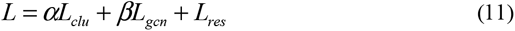

α, *β* and *θ* are used to balance the contribution of AE and GCN. In this study, we set α= 0.0001, *β* = 0.001 for all datasets. Finally, we choose the soft assignments in distribution Q as the final clustering results. The label assigned to cell ***i*** is:

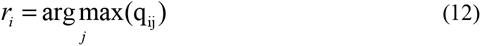

### Model construction for single-cell CCIs

Residual Network (ResNet) and GCN were used to build the interaction model of DeepCCI. The input of ResNet is the expression values of ligands in the source cell cluster and those of receptors in the target cell cluster. To fully utilize the information of each cell, we constructed the feature using the expression of the ligand of all cells in source cell cluster and the expression of the receptor of all cells in target cell cluster. To embed subunit information in deep learning, the geometric mean was applied to process the multi-subunit ligands or receptors present in each cell. To normalize feature data and improve model prediction speed, we use Singular Value Decomposition (SVD)^104^ method to reduce the feature data dimension before inputting the deep learning model. Simultaneously, the GCN model of this part receives the L-R pair graph that was constructed by the progressed scRNA-seq data and LRIDB. The nodes of L-R pair graph are the highly expressed L-R pairs in the scRNA-seq data, with the edges representing the relationships between these pairs. The highly expressed L-R pairs are defined based on the interaction probability. Specifically, we calculated the sum of the interaction probability of each L-R of all CCIs and selected the top 200 L-R pairs to construct the relationship between them. The correlation matrix **P** of all L-R pairs is built based on the Pearson correlation coefficient between each pair. Specifically, we use the threshold τ (default:0.95) to filter edges, and the adjacent matrix A^LR^ of L-R pairs can be written as:

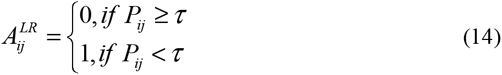

where 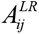 is the binary correlation matrix. In this way, we can capture the correlations between L-R pairs and explore these correlations to improve the classification performance for the interactions between two cell clusters. And then, the output of the ResNet and the GCN are applied by dot product for the following fully connected layer. The FC layers are used to predict L-R interactions with the cluster state. We assume that the ground-truth label of a significant interaction is **y**, where **y**_i_ denotes whether label **i** represent the significant interaction. The whole network is trained using the Focal loss^105^ as follows:

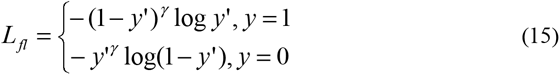

where *y*’ is the output of the FC layers activated by the sigmoid function.

### The evaluation criteria for DeepCCI

We compared the cell clustering results of DeepCCI with several scRNA-Seq analytical frameworks in terms of four clustering evaluation scores. ARI is used to compute similarities by considering all pairs of the cells that are assigned in clusters in the current and previous clustering adjusted by random permutation:

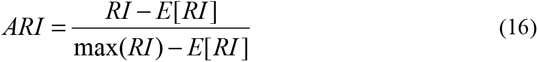

where the unadjusted rand index (RI) is defined as:

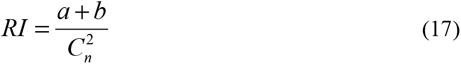

where *a* is the number of pairs correctly labeled in the same sets, and *b* is the number of pairs correctly labeled as not in the same data set. 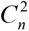 is the total number of possible pairs. E[RI] is the expected RI of random labeling.

The performance of DeepCCI on the prediction of CCIs was evaluated by using recall, precision, accuracy (ACC) and F1-score, which are well-known and have been widely used in the field of bioinformatics [51–55], defined as:

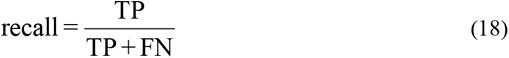

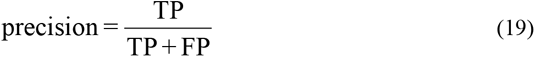

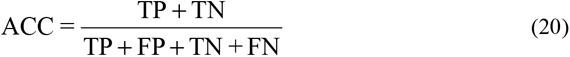

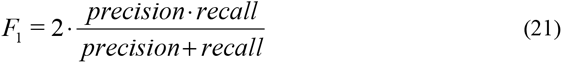

where TP represents true positive; TN, true negative; FP, false positive; FN, false negative.

Furthermore, the area under the curve (AUC) is an important metric of performance evaluation of the proposed model, defined as the area under the Receiver Operating Characteristic Curve (ROC) that can be used for visualizing the performance of interaction prediction.

### Method comparisons

We compare the performance of DeepCCI with seven other methods, including SingleCellSignalR, iTALK, CellPhoneDB, CellChat, CellCall, CytoTalk and NATMI. SingleCellSignalR uses a regularized expression product to compute ligand-receptor interaction scores and is the only tool reviewed that provides explicit cut-off values for this score to achieve appropriate false discovery rates based on empirical results. iTALK uses downstream analysis methods to curate the final list of significantly differentially expressed genes between cell clusters in single-cell RNA-seq data are identified, and these lists are analyzed for ligand-receptor pairs. CellCall and CytoTalk use known interactions between ligands, receptors and downstream targets to build a network of ligand-receptor relationships. Lists with multimeric proteins are used in CellPhoneDB and CellChat to assess whether all subunits are simultaneously expressed to identify likely functional ligand-receptor interactions. All the methods were applied to infer intercellular communications using their own default parameters.

### Availability of data and materials

The code for the DeepST algorithm, and a detailed tutorial are available at (https://github.com/JiangBioLab/DeepCCI).

The scRNA-seq data sets analyzed during the current study are publicly available. The benchmark data sets used for cell clustering can be downloaded from Gene Expression Omnibus (GEO) databases with accession numbers of GSE36552 (Yan), GSE75140 (Camp_Brain), GSE81252 (Camp_Liver), GSE85241 (Muraro), GSE94333(Adam), GSE109774 (QS Diaphragm, QS Limb Muscle, QS Lung, Qx Limb Muscle), GSE75688 (Chung); GSE65525 (Klein); GSE60361 (Zeisel). The Kolo data can be accessed from EMBL-EBI with an accession number of E-MTAB-2600. The Young data can be accessed from European Genome-phenome Archive (EGA) under study IDs EGAS00001002171, EGAS00001002486, EGAS00001002325, and EGAS00001002553. The labelled scRNA-Seq data for pancreatic islet are retrieved from GEO accession numbers GSE85241, E-MTAB-5061, GSE84133, GSE83139 and GSE81608, normalized with Seurat. The Human atopic dermatitis data can be downloaded from GEO database under accession code GSE147424 and the embryonic mouse skin datasets can be downloaded from GSE122043. The scRNA-seq data of human testicular cells were collected from GSE106487. The 10X Visium spatial transcriptomics dataset of coronal section of the mouse brain (https://support.10xgenomics.com/spatial-gene-expression/datasets) and seqFISH dataset of mouse organogenesis (https://content.cruk.cam.ac.uk/jmlab/SpatialMouseAtlas2020/) were obtained via the spatial single-cell analysis framework—Squidpy^106^(v1.1.0).

## Author’ contributions

Q.J. conceived and designed the study; Q.J., W.Y., Z.X., M.L. and C.X. performed the research; P.W., Y.C., J.Q., S.W. and R.C. collected and constructed the benchmark datasets; W.Y., Z.X., G.X, G.W. and Y.C. designed and implemented the computational framework with guidance from Q.J. H.N., P.W., and X.J. completed downstream analysis work. S.W., W.Z. and F.P. released the source code on GitHub; Q.J., W.Y., Z.X., Y.H., and M.L. wrote the paper with input from all other authors. All authors read and approved the manuscript.

## Funding

The National Natural Science Foundation of China (Nos. 62032007).

## Competing interests

The authors declare that they have no competing interests.

## Notes

### Competing Interest Statement

The authors have declared no competing interest.

